# Binding of SARS-CoV-2 fusion peptide to host membranes

**DOI:** 10.1101/2021.05.10.443474

**Authors:** Stefan L. Schaefer, Hendrik Jung, Gerhard Hummer

**Affiliations:** Department of Theoretical Biophysics, Max Planck Institute of Biophysics, 60438 Frankfurt am Main, Germany; Institute of Biophysics, Goethe University Frankfurt, 60438 Frankfurt am Main, Germany

## Abstract

During infection the SARS-CoV-2 virus fuses its viral envelope with cellular membranes of its human host. Initial contact with the host cell and membrane fusion are both mediated by the viral spike (S) protein. Proteolytic cleavage of S at the S2′ site exposes its 40 amino acid long fusion peptide (FP). Binding of the FP to the host membrane anchors the S2 domain of S in both the viral and the host membrane. The reorganization of S2 then pulls the two membranes together. Here we use molecular dynamics (MD) simulations to study the two core functions of the SARS-CoV-2 FP: to attach quickly to cellular membranes and to form an anchor strong enough to withstand the mechanical force during membrane fusion. In eight 10 *μ*s-long MD simulations of FP in proximity to endosomal and plasma membranes, we find that FP binds spontaneously to the membranes and that binding proceeds predominantly by insertion of two short amphipathic helices into the membrane interface. Connected via a flexible linker, the two helices can bind the membrane independently, yet binding of one promotes the binding of the other by tethering it close to the target membrane. By simulating mechanical pulling forces acting on the C-terminus of the FP we then show that the bound FP can bear forces up to 250 pN before detaching from the membrane. This detachment force is more than ten-fold higher than an estimate of the force required to pull host and viral membranes together for fusion. We identify a fully conserved disulfide bridge in the FP as a major factor for the high mechanical stability of the FP membrane anchor. We conclude, first, that the sequential binding of two short amphipathic helices allows the SARS-CoV-2 FP to insert quickly into the target membrane, before the virion is swept away after shedding the S1 domain connecting it to the host cell receptor. Second, we conclude that the double attachment and the conserved disulfide bridge establish the strong anchoring required for subsequent membrane fusion. Multiple distinct membrane-anchoring elements ensure high avidity and high mechanical strength of FP-membrane binding.

## Introduction

During infection, viruses first recognize and then enter their target cells. Coronaviruses such as SARS-CoV-2, the virus responsible for the ongoing COVID-19 pandemic, use their trimeric spike (S) glycoprotein for both tasks. The spike S1 subunit recognizes the human target cell by binding to the ACE2 receptor, and the S2 subunit then facilitates fusion of the viral membrane with host cellular membranes.^1–3^ To initiate fusion, in analogy to the hemagglutinin (HA) fusion protein of influenza, the SARS-CoV-2 S2 subunit is expected to first form one long trimeric coiled coil.^4^ This elongation would bring the fusion peptides (one per monomer) into the proximity of the membrane of the target cell. Binding to this membrane simultaneously anchors the S2 subunit in both the viral membrane (via its stalk) and the host membrane (via the FP). When the S2 subunit subsequently collapses to form a six-helix bundle in a proposed jack-knife mechanism, this pulls the host membrane and the viral membrane into proximity for eventual fusion.^4–8^

SARS-CoV-2 has two different routes of entry into the human host cell, either directly by fusion with the plasma membrane or by endosomal escape.^2,3,9^ In the latter pathway, the SARS-CoV-2 virion is endocytosed by the host cell after binding to the ACE2 receptor. As the membrane composition and pH of the endosome change, structural rearrangements may be induced in the S protein that facilitate membrane fusion. The virus then escapes the endosome before reaching the lysosome, releasing its RNA into the cytoplasm of the host.^9^

The FP of SARS-CoV-2 spike was identified as the 40 amino-acid long sequence just C-terminal of the S2′-cleavage site. ^2,10,11^ Upon proteolytic cleavage, S sheds its S1 subunit and releases the FP as the new N-terminus of its S2 subunit.^3,12,13^ Despite the concerted efforts to study the structure and flexibility of the S protein, ^8,14–18^ the structure of the FP after contact with the membrane has so far remained elusive. Nonetheless, mutagenic studies and electron spin resonance (ESR) experiments have provided some insight into the structure-function relationship of the SARS-CoV-1 FP. Using ESR, Lai et al. observed that both ends of the SARS-CoV-1 FP increased the order parameter of lipids that were spin labeled in their membrane interface region.^19^ N- and C-terminal fragments of the FP induced this effect individually; however, the intact FP showed the strongest effect on lipid order. Notably, no such ordering effect was observed after mutating the LLF motif in the N-terminal region of the FP to AAA. Mutation studies by Madu et al. confirmed the importance of the LLF motif for the fusion activity of the FP.^10^ Together, these experiments resulted in the idea of a bipartite fusion platform, with the LLF motif close to the N-terminus playing a crucial role. ^19^ The ability to increase the order parameter of spin labeled lipids was also confirmed for the SARS-CoV-2 FP.^20^

Given the lack of experimental structural data, we performed atomistic molecular dynamics simulations (MD) to elucidate the binding modes of the SARS-CoV-2 FP to the different host membranes it can encounter during infection. In a first set of simulations, we placed the FP in proximity to membranes mimicking the endosome and the outer leaflet of the plasma membrane, respectively. In this way, we could probe the spontaneous binding of the FP to these membranes. In a second set of simulations, we studied the mechanical strength of the FP membrane anchor. Starting with membrane-bound FP, we pulled the FP away from the membrane until it detached. Theoretical results suggest that in all binding modes observed here the FP anchoring is strong enough to support the complete fusion process. ^21^

## Results

### The FP in solution forms two short amphipathic helices

To explore the dynamics of the SARS-CoV-2 FP after S1 shedding and upon exposure to the surrounding medium, we performed MD simulations in aqueous solution, starting from the structure of the FP in intact S. Sequence and structural evidence suggests that the FPs of human infectious coronaviruses contain one highly conserved N-terminal amphipathic helix (NTH), a less conserved second amphipathic helix (AH2), and the C-terminal helix (CTH) (Figure 1a, c). The NTH is folded in the prefusion cryo-EM structure of the SARS-CoV-2 S protein resolved by Cai et al.^8^ During a 1 *μ*s simulation of the FP in water, the two C-terminal residues of the NTH (N824 and K825) quickly unfolded and the shorter NTH stabilized (Figure 1a, b). Formed by consecutive residues, the LLF motif is spread across both faces of the NTH, as the two leucines are part of the predicted hydrophobic face and the phenylalanine is not. At the C-terminal end of the FP segment, the EM structure of S shows three helical segments interrupted by short disordered regions. In our simulation of FP in aqueous solution, the first segment expanded to the beginning of the second segment to form a single contiguous helix (AH2), the remainder of the second segment unfolded, and the third segment retained its helical structure. These two distinct short helices (AH2 and CTH) are connected via a short loop. Whereas the amino-acid sequence of CTH is highly conserved and shows no strong amphipathic properties, AH2 is less conserved but carries a strong hydrophobic moment (Figure 1c). Notably, AH2 and the CTH are additionally connected via a fully conserved disulfide-bridge.

**Figure 1:**
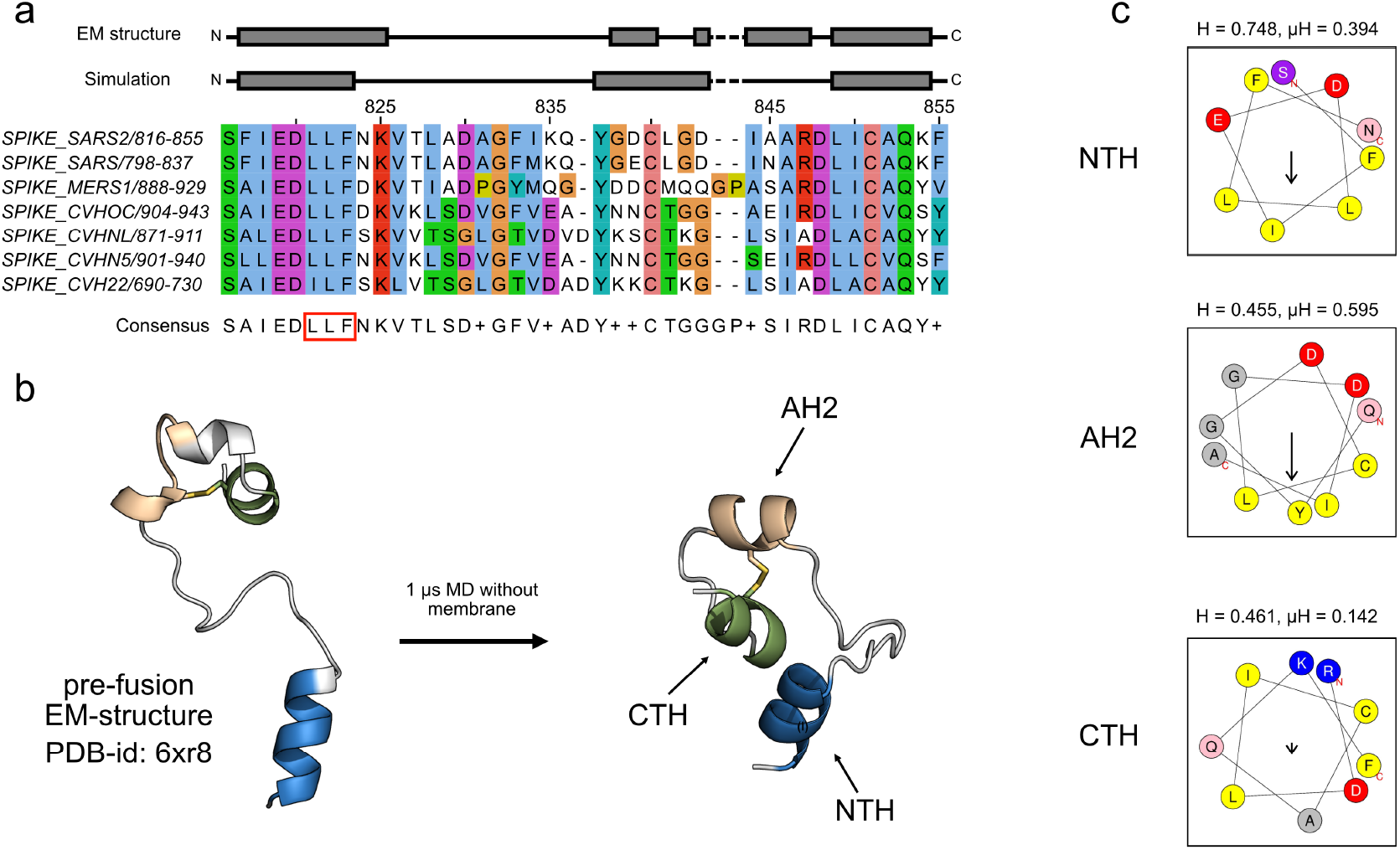
Amphipathic helices in SARS-CoV-2 fusion peptide. (a) Alignment of the FP region of human infectious coronaviruses and helix assignment from the cryo-EM structure (PDB ID: 6XR8, top)^8^ and from our simulation of the peptide without a membrane present (bottom). The red box marks the LLF motif in the N-terminal region. The fully conserved cysteines are at positions 840 and 851. The alignment was calculated using Clustal Omega. ^22^ See SI Figure S1 for a larger alignment that includes other betacoronaviruses. (b) Structural change of the FP in a 1 *μ*s simulation in water and NaCl in cartoon representation (NTH in blue, AH2 in beige, CTH in green). (c) Amphipathic profiles of the NTH (top), the AH2 (middle) and the CTH (bottom) from SARS CoV-2 with hydrophobicity H and hydrophobic moment *μ*H calculated with HeliQuest. ^23^

### NTH binds membranes with its amphipathic face

We reasoned that the two amphipathic helices NTH and AH2 may insert into the human membranes to anchor the S protein for membrane fusion. To test this hypothesis, we performed MD simulations of FPs placed near lipid bilayers. In eight independent MD simulations of 10 *μ*s each, we observed five spontaneous insertion events of the NTH into the membrane interface. Four of these events occurred on the endosomal membrane, meaning that all replica simulations with this composition ended with the NTH inserted (Table S1). In three cases the NTH was the part of the peptide that first created a stable contact. Two other spontaneous insertion events were mediated by membrane contacts of AH2 and CTH. Notably, once the NTH bound to the membrane, it remained bound for the entire duration of the simulations. Only in one of the eight simulations did the FP not stably insert into the membrane.

In all three cases in which the NTH established the first stable contact, the binding followed a consistent path. First, F817 penetrated below the phosphate headgroup region of the membrane, after which the rest of the NTH bound the membrane with its predicted hydrophobic face. NTH binding stabilized in two slightly different ways: In three simulations (Figure 2; runs 1, 2, 4), F823 flipped its orientation after being bound to the endosomal membrane for ≈ 0.7, 2, and 3 *μ*s, respectively, so that its aromatic sidechain became completely buried under the lipid headgroup region. This led to an overall deeper insertion of the NTH, where all three residues of the LLF motif became deeply burrowed into the membrane. We found that, once flipped, F823 can transition between a favored deep insertion state and a shallow state (Figure S2). In the remaining insertion event into the endosomal membrane, the NTH established stable binding without F823 flipping in the simulated time and hence the helix remained on top of the membrane interface. In this shallower binding state, only the residues of the predicted hydrophobic face of the NTH inserted into the headgroup region (Figure 1c), as did the disordered region immediately downstream of the NTH (V826–A829).

**Figure 2:**
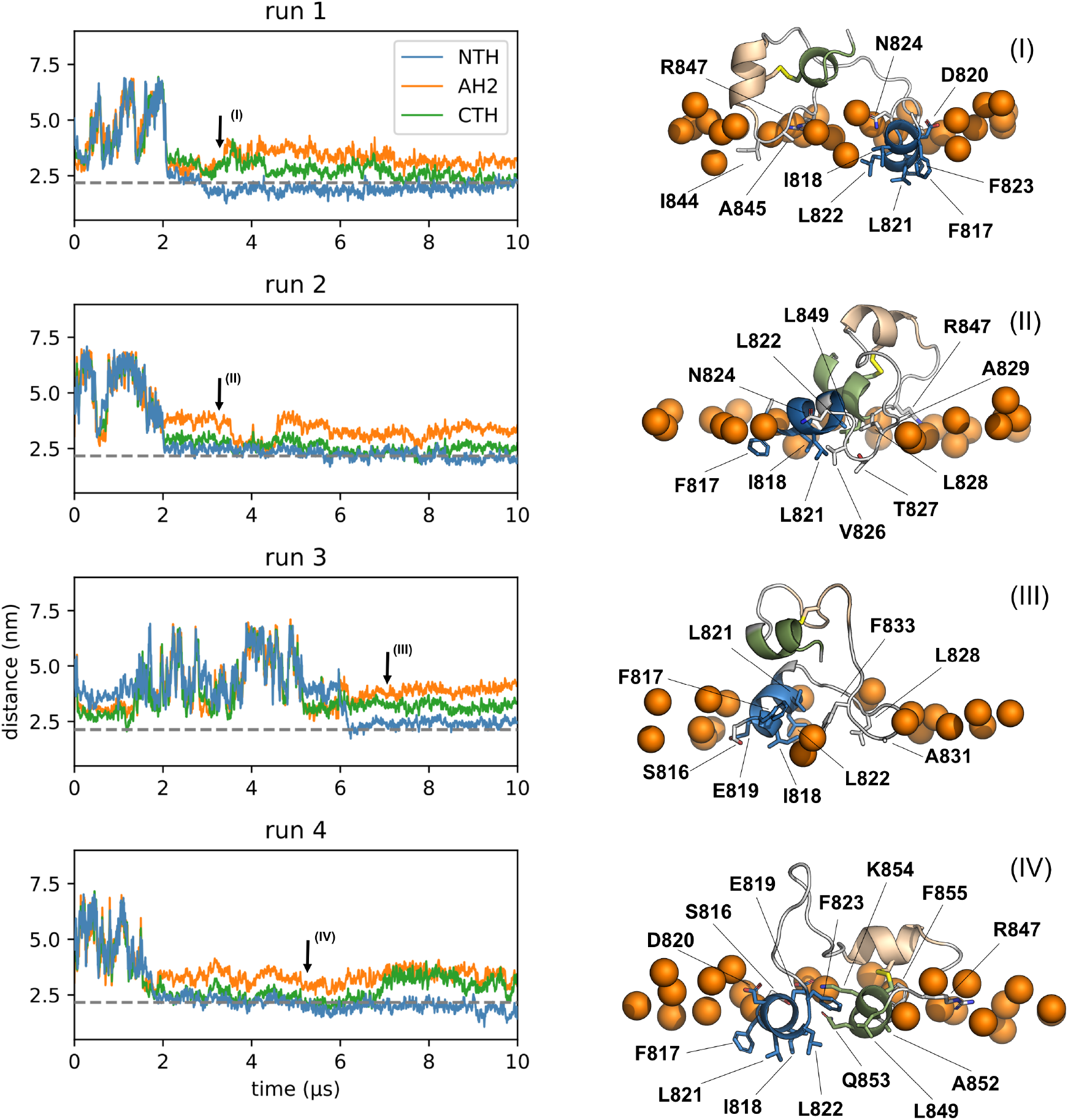
NTH binding to the endosomal membrane. (Left) Distances of the centers of mass of the three helices and the center of mass of the membrane. The average phosphate position of the bound leaflet is indicated by a gray dotted line. Arrows indicate times for snapshots in right panel. (Right) Representative snapshots of states bound to the membrane. Colors as in Figure 1 (NTH: blue, AH2: beige, CTH: green). Membrane-inserted residues are labeled and shown in licorice representation. The upper membrane boundary is indicated by phosphate headgroups of nearby lipids (orange spheres).

In the simulations with the mimetic of the outer leaflet of the plasma membrane, we observed one spontaneous NTH binding event, after the CTH and AH2 had already been inserted for more than 5 *μ*s (Figure 3, run 4). All three helices stayed bound to the membrane for ≈ 0.5 *μ*s until the NTH lost membrane contact. To relieve the lateral-pressure asymmetry between the two leaflets caused by inserting a large structure into only the top leaflet of a finite-size membrane patch, we selected this relatively short lived state with all three helices bound, removed 5 lipid molecules of the over-compressed leaflet and restarted the simulation. In this pressure-relieved mode, all three helices remained stably bound for the entire simulated time (>3 *μ*s, Figure S3a). F823 flipped into the membrane under the phosphate headgroups after ≈ 1.3 *μ*s, which led to deeper NTH insertion.

**Figure 4:**
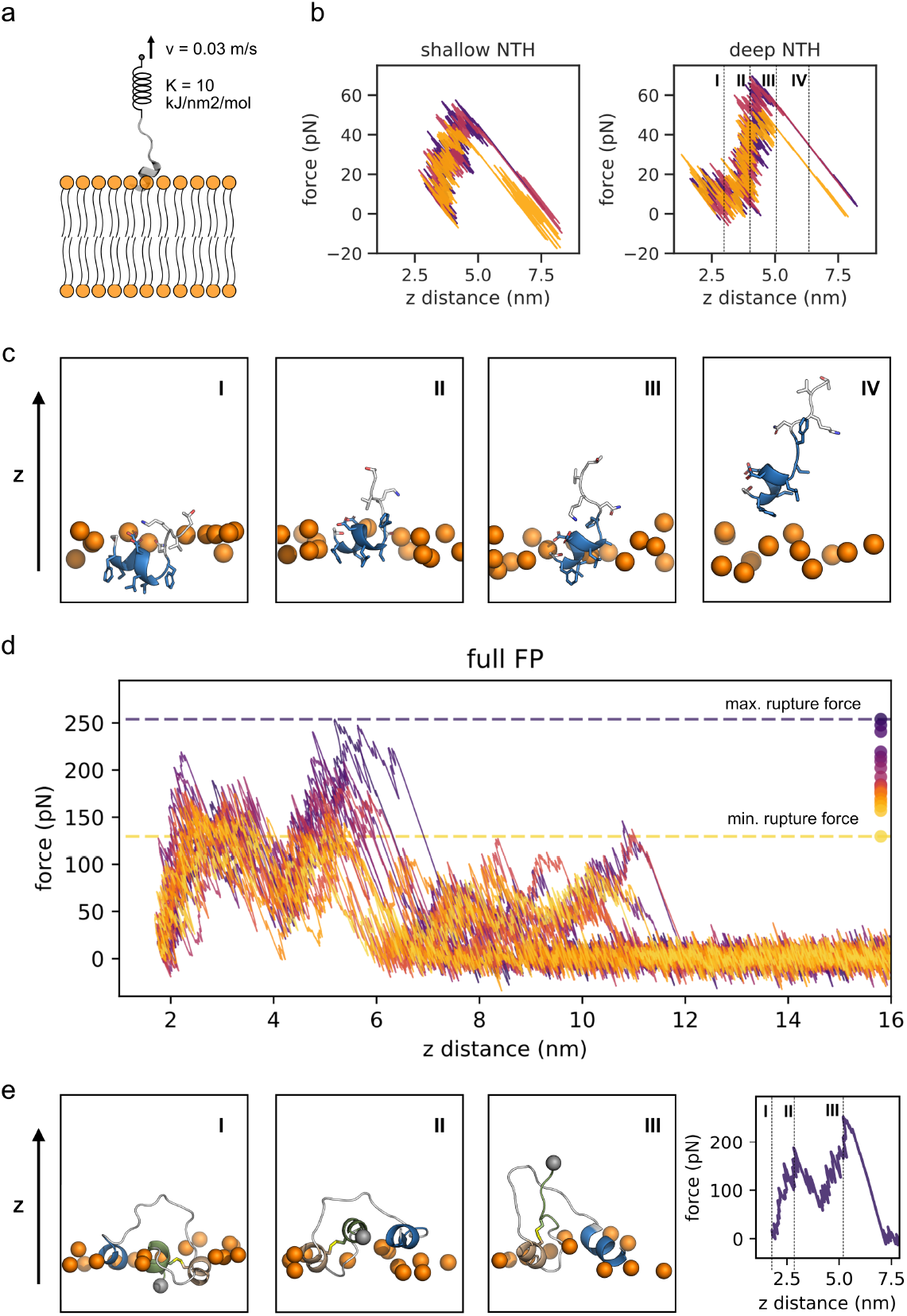
Mechanical strength of membrane anchoring by FP. (a) Schematic of pulling on the FP mediated by a spring whose end point is moving at a constant velocity. (b) Force-extension curves for pulling the bound NTH off the endosomal membrane. The x-axis shows the distance between the NTH center of mass and the membrane center of mass only in z-direction. The initial structures were taken from one simulation with the endosomal membrane before and after F823 flipping. The dashed lines in the right panel indicate frames for the snapshots of one replica (purple line) depicted in c. The coloring of the NTH is according to Figure 1. The upper membrane boundary is indicated by phosphate headgroups of nearby lipids (orange spheres). (d) Force-extension curves for pulling the full length FP with all three helices inserted via its C-terminus off the outer plasma membrane. Individual trajectories are colored according to their peak rupture force, as indicated by the points on the right and dashed lines for the overall minimum and maximum. Axes as in b. (d) Snapshots of the replica that resulted in the highest rupture force at distances indicated by the dashed lines in the corresponding force-extension curve on the right. Coloring according to Figure 1. The C-terminus is depicted as a gray sphere.

### The C-terminus of the FP binds via flexible elements

In all our simulations, we observed membrane binding also with the C-terminal end of the FP. Binding involved AH2, the CTH and flexible elements flanking AH2 at both ends. In some cases, these membrane interactions were relatively short lived (see, e.g., Figure 2, run 3), in other cases binding was stable for more than 6 *μ*s. Interestingly, some stable interactions were mediated by only few but tight interactions (see e.g., Figure 3, run 2), whereas other more extensive interactions were short-lived. The short hydrophobic stretches that inserted most frequently are centered around residues I834, L841 and I844. As these residues are located at the borders of the AH2 and the CTH, their binding repeatedly led to the insertion also of residues of the respective neighboring helix. This coupled insertion was especially common in the case of AH2, which, by this process, was guided into the membrane interface with its predicted hydrophobic face. Notably however, the short AH2 and to an even larger extent the CTH, in some cases partially unfolded to flexible amphipathic structures when bound to the membranes (Figure S4).

**Figure 3:**
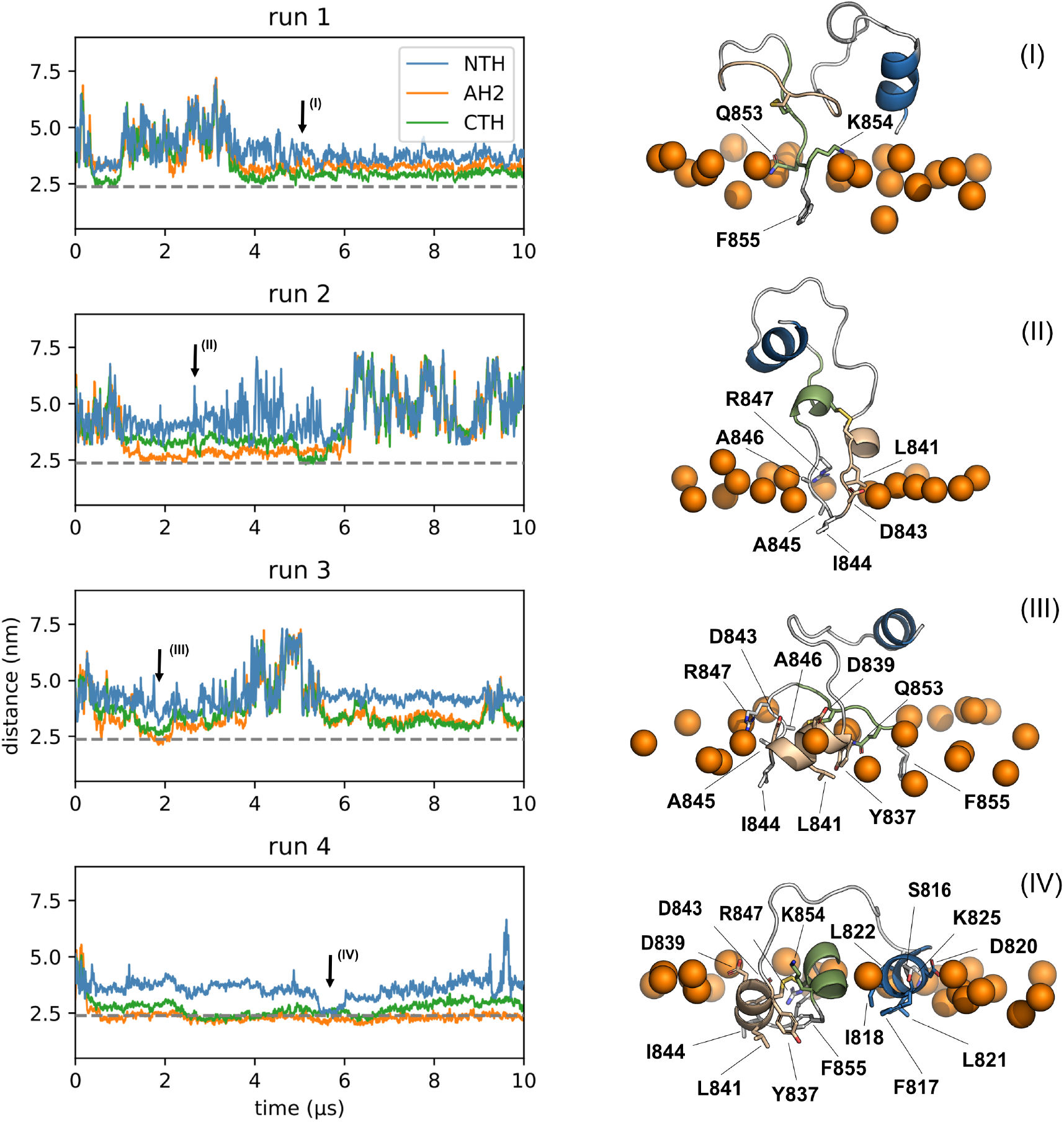
FP binds outer plasma membrane mimetic in different modes. (Left) Distances between the centers of mass of the three helices and the center of mass of the membrane. The average phosphate position of the bound leaflet is indicated by a gray dotted line. Arrows indicate times for snapshots in right panel. (Right) Representative snapshots of states bound to the membrane. Colors as in Figure 1 (NTH: blue, AH2: beige, CTH: green). Membrane-inserted residues are labeled and shown in licorice representation. The upper membrane boundary is indicated by phosphate headgroups of nearby lipids (orange spheres).

### The inserted NTH can withstand high pulling forces

We determined the strength of the membrane anchoring by subjecting the the C-terminus of the FP to mechanical force. This process mimics the forces experienced by the FP during its presumed primary function of pulling host and viral membrane into proximity. In our simulations, we applied force to the C-terminus of the FP in a direction normal to the membrane. By pulling the C-terminal end up via a harmonic spring moving at constant velocity, the force applied to the peptide increases more or less linearly in time, until peptide segments, and ultimately the entire peptide, are pulled out of the membrane. Each of these events results in a distinct drop in force. By pulling the bound NTH out of the endosomal membrane, we found that the binding of the NTH alone can withstand pulling forces between 40 and 65 pN. As shown in Figures 4b and c, the applied force increased as long as the peptide was still in contact with the membrane. When the peptide lost its last contact with the membrane, it detached in a sudden transition. Higher forces were needed to pull the NTH out of the deeply bound state with inserted F823 (Figure 4b, right). Interestingly, even though F823 was pulled out of the membrane along this path (Figure 4c, II), the shallow state did not appear as a distinct intermediate in the pulling traces. Nonetheless, the forces required to completely detach the NTH from the deep state are 10-15 pN higher compared to the shallow binding mode.

### Additional insertion of AH2 and CTH dramatically stabilize membrane anchoring

We performed additional pulling simulations with the full length FP, starting from a binding mode with all three helices inserted into the membrane interface. The initial structure for the 20 replica pulling simulations was taken from an unbiased simulation of FP on the outer plasma membrane (for details see methods). Having all three helices inserted increased the mechanical stability of the FP membrane anchoring. With rupture forces ranging from 130 to 254 pN, these simulations reveal that binding of the full FP is about two to three times as strong as binding of the NTH alone (Figure 4d, dashed lines and dots). Additional pulling simulations of structures of the same simulation at different times show similarly high rupture forces (Figure S3b).

For the full-length FP, the force-extension curves also changed in character compared to the NTH alone (Figures 4b and d). We consistently observed multiple distinct rupture peaks. The first two of these peaks were observed in all 20 replicas. The events resulting in the two force peaks are connected to each other because the force does not drop back to zero in-between them. One might assume that these peaks were the result of the CTH and AH2 detaching individually. However, visual inspection revealed that this is not the case. Instead, the first peak corresponds to the extraction of the neutral F855 terminus. In most cases, this is closely associated with the detachment of Y837 and L841 of the AH2 from the membrane, after which AH2 stands upright. This causes a sudden gain in flexibility of the C-terminus that results in a local minimum (above 4 nm) in the force-extension curve (Figure 4d and e, Movie S5). However, at this point, I844 in the short loop between the AH2 and the CTH still remains inserted in all simulations. The detachment of I844 then results in the second rupture peak. In 35% of the simulations the entire peptide was detached from the membrane after the second force peak, meaning that all three helices detached nearly simultaneously with the detachment of I844. In the remaining 65% of pulls, parts of the long linker and the NTH stayed bound independently of the AH2 and the CTH and only detached later. This then resulted in a series of additional force peaks, with rupture forces in the same regime as for the isolated NTH fragment.

## Discussion

### Two separate binding regions increase the likelihood to stay bound under load

Due to the connection via a long disordered linker, the NTH and the two more C-terminal helices, AH2 and CTH, act relatively independent from one another. Nonetheless, binding of one part to the membrane automatically places the other part close to the membrane as well, thus making it easier for it to bind. Therefore, their binding behaves cooperative while at the same time unbinding of one part does not necessarily lead to rapid dissociation of the FP away from the membrane. Hence, we hypothesize that the architecture of the FP results in a fast *k*_on_-rate, thanks to the small amphipathic insertion elements, and a slow *k*_off_-rate thanks to multiple interaction elements and their geometric arrangement. For SARS-CoV-2, the concept of avidity – with multiple spread-out interactions maintaining a bound state – has emerged at multiple levels: in the ACE2-S interaction,^24^ in the virion-host interaction,^14^ and here in the FP-membrane interaction.

A particularly interesting force-bearing element in the FP is the disulfide bridge connecting the centers of AH2 and the CTH. In the lead-up to membrane fusion and thus viral infection, we expect force to act on the C-terminus of the FP, as in our MD simulations. By directing this force to the center of a membrane-anchored AH2 via the covalent disulfide bridge, instead of applying it to the C-terminus of the helix, AH2 has to be pulled out of the membrane all at once instead of lifting it from the end. With this in mind, it is therefore not surprising the two cysteines are fully conserved in the betacoronavirus family (Figure S1). In this way, AH2 can be kept small in footprint for rapid insertion yet sustain strong binding also under significant force.

The behavior we observed is in perfect accordance with the bipartite fusion platform idea as proposed by Lai et al.^19^ They measured that both ends of the FP of SARS-CoV-1 could individually increase the membrane order; however, when covalently bound they observed the strongest effect. This underlines our idea of the two sides of the FP acting cooperatively, promoting each other’s membrane binding and stabilizing the anchoring overall. In addition, we can directly link the described crucial role of the LLF motif to our observed NTH binding as it consistently includes the insertion of both leucines (L821 and L822) into the glycerol backbone region of the membrane lipids. Remarkably, we found that F823, despite being placed on the hydrophilic face of the amphipathic helix, flipped its orientation so that its sidechain became membrane inserted in the deeply bound state of the NTH. As we showed, the deep binding state associated with F823 insertion increases the pulling force that the NTH can withstand, which in turn supports the fusion activity of the FP.

Two other MD simulation studies recently addressed membrane binding by the FP.^25,26^ Gorgun et al. used a truncated version of the FP where they cut it behind L841, right where we observe the AH2. In a series of 30 short (300 ns) simulations, they observed binding of the FP to a highly mobile membrane mimetic (HMMM) membrane model. They identified three binding modes: one with the loop inserted into the membrane, one with the NTH inserted and a last binding mode where the whole FP acquires helical structure and inserts on top of the membrane. ^25^ The first two of these binding modes stand in good agreement with our observations. However, it is worth noting that in the case of the long loop binding, we often only observe shallow insertion of the involved residues. Our comparably long trajectories additionally reveal that those loop insertions may play a role in guiding the larger helical parts into the membrane, but do not remain stably bound on their own. The third binding mode reported by Gorgun et al. ^25^ was not observed in our simulations. Nevertheless, this does not mean that it may not realistically occur also with the full-length FP, if simulated longer. The second study, by Khelashvili et al.,^26^ focuses on the role of Ca^2+^-ions for FP binding. In a large set of 1 *μ*s long simulations, they also identified the NTH and the AH2 as predominantly helical and confirm the binding of the NTH region to the membrane as the most prevalent binding mode in their Ca^2+^ coordinated structures.^26^

### Differences in lipid density may alter preferred binding mode

The binding modes we described seem to loosely group into the NTH binding to the endosomal membrane and C-terminal regions binding to the outer plasma membrane mimetic. The most pronounced difference between the two membranes is their density and in particular the density in the lipid headgroup region. The outer plasma membrane has a higher content of cholesterol and sphingolipids, which pack tightly together, thus increasing the overall lipid density. Adding to this, the outer plasma membrane contains relatively fewer lipids with small headgroups, which would lower the pressure in the interface region. Specifically, in the endosomal membrane, the two abundant lipids POPE and BMP decrease the density in this area. Therefore, it is not surprising that the stably folded NTH can relatively easily insert into the endosomal membrane, but only does so rarely with the outer plasma membrane. By contrast, the high density of the outer plasma membrane may favor the insertion of smaller, often disordered hydrophobic stretches such as the ones around I844. It is therefore tempting to speculate that the observed differences in insertion behavior may indeed be representative of the initial binding of FPs into the comparably soft and compressible endosomal membrane and the rigid and dense plasma membrane, both of which have been reported to be targeted by SARS-CoV-2.^2,3,9^

### Finite size effects may impair binding of the whole FP

In MD simulations of membrane systems, the spontaneous binding of a peptide into the membrane is artificially hindered. Binding to only one leaflet of a small membrane patch inevitably increases the lateral pressure in that leaflet and creates a significant asymmetry in the packing of lipids in the two leaflets. This creates an artificial energetic penalty that competes with the binding free energy of the amphipathic peptide. Unfortunately, this effect is difficult to correct for, short of performing simulations with prohibitively large boxes or with preemptively removed lipids from one leaflet. Therefore, we expect that in our simulation setup binding is weakened. We speculate that we would have seen more and longer-lived binding modes to membranes with asymmetric leaflet density. Furthermore, the lower penalty from the lateral pressure would have likely led to the simultaneous binding of two or all three helices. This is underlined by the fact that in the initial, symmetric simulations the binding of all three helices together is only transiently stable, whereas relieving the pressure in the bound leaflet makes the same binding mode stable for much longer. However, taken together with the fact that, despite the energetic penalty, this state occurred at all, we hypothesize that it may represent the energetically most favorable binding mode of the FP.

### Binding of few FPs may be strong enough to facilitate membrane fusion

Kozlov and Chernomordik made a theoretical estimate of the forces acting on an influenza HA FP during the fusion process. ^21^ During fusion, the HA2 subunit folds back onto itself and creates a six helix bundle.^4–6^ They estimated that the energy released from this folding would give rise to ≈ 8 pN of pulling force acting on each of the three FPs of HA. For SARS-CoV-2, the fusion process is thought to be similar to that mediated by HA, and we therefore expect that the forces are also comparable.^4^ The 130-250 pN forces required to detach the bound FP in our simulations greatly exceed the ones necessary according to these theoretical considerations. Here we emphasize that the time scale of 100 ns to 1 *μ*s over which the force is ramped up in the simulations is in a range not unreasonable for the spike refolding from prefusion to postfusion conformations. Even if at a lower force loading rate dissociation happened already at a somewhat lower force, that force would still likely exceed substantially the force required for fusion. ^21^ Whereas already the bound NTH alone can sustain such forces, the full FP is anchored even more strongly by its three helices. As discussed, the fully conserved cysteine-bridge emerged as an important mechanical stabilizer. The disulfide bond connects the AH2 and the CTH at their centers, and thus directs the pulling force away from the ends of the AH2. Instead of a sequential detachment of single amino acids, each event with a comparably low energetic barrier, the application of force to the center of the helix favors a pathway with a high barrier in which the entire hydrophobic face of the AH2 is detached at once, before then I844 is pulled out normal to the membrane (Figure 4e). The structure with a disulfide bridge thus endows the FP with high anchoring strength that is reminiscent of the catch bonds giving cell-cell contacts high mechanostability.^27^

The observed stability of the binding raises the question of how many bound FPs are required to be engaged for successful fusion. With all three of its helices bound, the full estimated pulling force could be borne by just one FP. ^21^ This may ultimately increase the infection success of the virus, as it would reduce one source of failure.

## Conclusions

From atomistic molecular dynamics simulations, we gained a detailed view of the interactions between the SARS-CoV-2 FP with lipid bilayers mimicking the endosomal membrane and the outer leaflet of the plasma membrane. In our MD simulations, we observed multiple spontaneous membrane insertion events. In all four runs with the more flexible and less packed endosomal membrane, the FP eventually bound into the lipid bilayer with its NTH. Adhered to membranes, the FP retained much of the secondary structure seen in pre-fusion spike. The FP folds such that two short amphipathic helices can bind to the membrane interface with well defined hydrophobic faces. Additional highly flexible hydrophobic stretches can prime the membrane insertion process and stabilize the bound state. The NTH and the two C-terminal helices AH2 and CTH are separated by a flexible linker and can therefore insert independently. Insertion of one, however, likely promotes the insertion of the other, simply by disallowing escape away from the membrane. Insertion of all three helices at the same time was observed rarely in our simulations. Nonetheless, we could show that by relieving lateral pressure in the exposed leaflet, we could stabilize a binding mode with all three helices inserted fully. We therefore expect that FP binding will eventually converge to all three helices bound to the membrane in the course of a real infection event.

We could show that the bound FP — even though it is bound only to the interface of the membrane — can withstand large pulling forces exceeding 200 pN. In fact, the forces are so high that binding of only one of the three FPs of S may suffice for membrane fusion.^21^ The strong anchoring force hints at a connection of the architecture of the FP and the infection success of the SARS-CoV-2 virus. The Cys-Cys disulfide bond linking the centers of AH2 and CTH emerged as an important stabilizer. By transmitting the force load during the membrane fusion process to the center of the membrane anchored AH2 instead of its C-terminus, AH2 has to be pulled out of the membrane all at once to detach the FP from the membrane. We speculate that by spreading the membrane interaction across multiple distinct elements, with NTH, AH2, CTH and the intervening amphipathic loops all connecting to the membrane, the virus achieves a trade-off between rapid insertion of individually small elements into the membrane and their firm membrane anchoring.

The design principles for the SARS-CoV-2 fusion peptide emerging from our MD simulations could be relevant, on the one hand, for the design of fusion inhibitors and, on the other hand, for the biotechnological development of membrane anchors and fusogens, e.g., for drug delivery applications.

## Materials and Methods

### General simulation parameters

All MD simulations were performed with Gromacs 201 8.8^28^ using the TIP3P water model and the CHARMM36m forcefield. ^29^

### FP in water

The FP was extracted from the S protein prefusion cryo-EM structure (PDB ID: 6XR8) and used as the input for the CHARMM-GUI solution builder.^8,30^ Its C-terminal end was modeled as a methylamidated C-terminus, to account for the continuation of the peptides. The disulfide-bridge between Cys840 and Cys851 was added. The peptide was solvated with TIP3P water and 0.15 M NaCl.

Energy minimization was done using a steepest descent algorithm for 5000 steps with restraints as described in table S4. Subsequently, the system was equilibrated for 125 ps, with a timestep of 1 fs and Nosé-Hoover temperature coupling^31,32^ with a reference temperature of 310.15K and the same restraints as used during the minimization.

Finally, 1 *μ*s of production simulation was performed. For this, the timestep was increased to 2 fs and temperature coupling was handled by the velocity-rescale algorithm^33^ coupled to the protein and the rest of the system individually. Isotropic pressure coupling was handled by the Parrinello-Rahman algorithm with compressibility *K_xyz_* = 4.5 × 10^-5^ bar^-1^.^34^

### Amphipathic helix prediction

To analyze the physicochemical properties of the FP and in particular its amphipathic helices we used the HeliQuest web-server.^23^ Due to how short the AH2 and the CTH are, 1-2 neighboring residues were included on both sides of the observed helices.

### Membrane compositions

To get insight into both membranes the FP can come into contact with in a realistic infection case, we built systems with two different membrane models. One is a recreation of the the outer leaflet plasma membrane used by Lorent et al. ^35^ and is characterized by a high amount of cholesterol and sphingolipids. These two types of lipids are known to pack tightly together creating a relatively stiff membrane. Moreover, this membrane has a high phosphatidylcholine (PC) and a low phosphatidylethanolamine (PE) content.

The second, the endosomal membrane model, was built based on results reviewed by van Meer et al. ^36,37^ Notably, this membrane includes the late endosome specific lipid BMP and lipids with small, negatively charged headgroups.

The detailed membrane compositions are summarized in tables S2 and S3.

### Membrane simulations

The unbiased simulations of the fusion peptide on the different membranes were set up using CHARMM-GUI membrane builder,^30,38^ with TIP3P water and 0.15 M NaCl. In both cases the FP was placed in close proximity to the membrane interface, but not bound to it (shortest atom-atom distance ≈ 5 Å). The membranes for the plasma membrane and the endosomal membrane systems were built symmetrically with 150 and 160 lipids per leaflet respectively. All simulations were minimized for 5000 steps using steepest descent and restraints as described by table S5. Subsequently, both systems were equilibrated in six equilibration steps with decreasing restraints (table S5). During equilibration, the temperature coupling was handled by the Berendsen thermostat^39^ with a reference temperature of 310.15 K. Additionally, semiisotropic Berendsen pressure coupling^39^ with reference pressure of 1 bar and compressibility *K_z_* = *K_xy_* = 4.5 × 10^-5^ bar^-1^ was applied for all but the first two equilibration rounds. After equilibration, the production runs were started without restraints. For the production simulations, temperature coupling was handled by the velocity rescale algorithm^33^ coupled to the water together with the ions, the peptide and the membrane individually with a coupling constant of one picosecond. Semiisotropic pressure coupling was handled by the Parrinello-Rahman algorithm^34^ with a reference pressure of 1 bar and compressibility *K_z_* = *K_xy_* = 4.5 × 10^-5^ bar^-1^. Pressure coupling was applied every 5 ps. To create independent replicas, all four production runs of both systems were initialized with different random velocities generated according to the Maxwell-Boltzmann distribution.

### Constant velocity pulling

For constant velocity pulling of the NTH, two structures from one of the endosomal membrane simulations (run 1) were used as starting structures — one at 2.5 *μ*s and the other at 3.5 *μ*s. To be able to isolate the strength of the NTH binding from effects caused by the rest of the FP, the peptides were then shortened to T827 and the new C-terminal end was again methylamidated. In addition, the box size was increased in the z-direction to 18.2 nm. To do so, all water molecules and ions were removed after increasing the box size, and the systems were re-solvated and re-ionized with 0.15M NaCl. Subsequently, the solvent was equilibrated for 1 ns, while heavily restraining the peptide and the lipid positions (same restraints as for 1^st^ equilibration; table S5). Replica simulations were started from the same initial structure but with randomized initial velocities generated according to the Maxwell-Boltzmann distribution.

The same approach was used for pulling the FP with all three helices bound off the membrane. There, the structure of the plasma membrane simulation run 4 at 5.7 *μ*s (structure in figure 3, bottom) was used. In contrast to the NTH pulling simulations, the peptide was not shortened and the box size had to be increased in *z*-direction to 35 nm. In addition, the lateral pressure of the bound leaflet was relieved by removing 5 lipids from it (2 cholesterol, 1 PLPC, 1 PSM, 1 NSM). Binding modes for simulated pulling were taken at 0, 1, 2, and 3 *μ*s of an unbiased simulation with said system.

For the simulations, the Gromacs 2018.8 pull code^28^ was used to move the C-terminal carbon atom, away from the membrane center of mass at a constant velocity of 0.03 ms^-1^. A spring with a weak force constant of 10 kJ mol^-1^ nm^-2^ was chosen to connect the dummy and the C-terminal carbon. The pulling force was only applied in the z-direction and the simulations ended, when the distance between the center of mass of the membrane and the C-terminal carbon was more than 0.49 times the box size in *z.*

## Notes

### Competing Interest Statement

The authors have declared no competing interest.

