## Supporting Information for "Binding of SARS-CoV-2 fusion peptide to host membranes"

### Supporting Information Available

### Tables

Table S1: FP secondary-structure elements in membrane contact during initial membrane binding event that led to stable binding and in final trajectory frame (initial/final).

| membrane system | run no. | NTH bound | AH2 bound | CTH bound |
|---|---|---|---|---|
| endosomal | 1 | ✓/✓ | ✓/✓ | ✗/✓ |
| endosomal | 2 | ✓/✓ | ✗/✗ | ✓/✓ |
| endosomal | 3 | ✗/✓ | ✓/✗ | ✓/✗ |
| endosomal | 4 | ✓/✓ | ✗/✗ | ✓/✗ |
| outer plasma | 1 | ✗/✗ | ✗/✓ | ✓/✓ |
| outer plasma | 2 | ✗/✗ | ✗/✗ | ✗/✗ |
| outer plasma | 3 | ✗/✗ | ✓/✓ | ✓/✗ |
| outer plasma | 4 | ✗/✗ | ✓/✓ | ✓/✓ |

Table S2: Outer plasma membrane-like lipid membrane composition as created by Lorent et al.[35]

| Lipid | Acyl chains | Full name | Abundance % |
|---|---|---|---|
| CHOL | | Cholesterol | 40 |
| PSM | 18:1/16:0 | N-palmitoyl-D-erythro-sphingosylphosphorylcholine | 12 |
| NSM | 18:1/24:1 | N-nervonoyl-D-oleoyl-sphingosylphosphorylcholine | 9.3 |
| LSM | 18:1/24:0 | N-lignoceroyl-D-oleoyl-sphingosylphosphorylcholine | 8 |
| PLPC | 16:0/18:2 | 1-palmitoyl-2-linoleoyl-sn-glycero-3-phosphocholine | 14.7 |
| SOPC | 18:0/18:1 | 1-stearoyl-2-oleoylphosphatidylcholine | 6.7 |
| PAPC | 16:0/20:4 | 1-palmitoyl-2-arachidonoyl-glycero-3-phosphocholine | 5.3 |
| PLA20(PE) | 18:0/20:4 | 1-O-stearoyl-2-O-arachidonoyl-glycero-3-phosphoethanolamine | 2.7 |
| SAPS | 18:0/20:4 | 1-stearoyl-2-arachidonoyl-glycero-3-phosphoserine | 1.3 |

Table S3: Late endosomal membrane composition

| Lipid | Acyl chains | Full name | Abundance % |
|---|---|---|---|
| CHOL | | Cholesterol | 27 |
| PLPC | 16:0/18:2 | 1-palmitoyl-2-linoleoyl-sn-glycero-3-phosphocholine | 22 |
| POPE | 16:0/18:1 | 1-palmitoyl-2-oleoyl-glycero-3-phosphoethanolamine | 16 |
| BMP | 18:1/18:1 | Bis(monoacylglycero)phosphate | 11.6 |
| POPC | 16:0/18:1 | 1-palmitoyl-2-oleoyl-glycero-3-phosphocholine | 11 |
| SSM | 18:1/18:0 | N-stearoyl-D-erythro-sphingosylphosphorylcholine | 7.3 |
| SAPI | 18:0/20:4 | 1-stearoyl-2-arachidonoyl-sn-glycero-3-phosphoinositol | 3.65 |
| SAPS | 18:0/20:4 | 1-stearoyl-2-arachidonoyl-sn-glycero-3-phospho-L-serine | 1.45 |

Table S4: Restraints used during energy minimization and equilibration of the FP in aqueous solution. Values in units of $\mathrm{kJ\,mol^{-1}\,nm^{-2}}$

| | **Backbone** | **Sidechains** | **Dihedrals** |
|---|---|---|---|
| EM | 400 | 40 | 4 |
| EQ | 400 | 400 | 4 |

Table S5: Restraints used during energy minimization and equilibration of the endosomal and outer plasma membrane systems in $\mathrm{kJ\,mol^{-1}\,nm^{-2}}$

| | **Time [ns]** | **Timestep [fs]** | **Backbone** | **Sidechains** | **Lipids** | **Dihedrals** |
|---|---|---|---|---|---|---|
| EM | | | 4000 | 2000 | 1000 | 1000 |
| $1^{st}$ EQ | 0.125 | 1 | 4000 | 2000 | 1000 | 1000 |
| $2^{nd}$ EQ | 0.125 | 1 | 2000 | 1000 | 400 | 400 |
| $3^{rd}$ EQ | 0.125 | 1 | 1000 | 500 | 400 | 200 |
| $4^{th}$ EQ | 0.5 | 2 | 500 | 200 | 200 | 200 |
| $5^{th}$ EQ | 0.5 | 2 | 200 | 50 | 40 | 100 |
| $6^{th}$ EQ | 0.5 | 2 | 50 | 0 | 0 | 0 |

### Figures

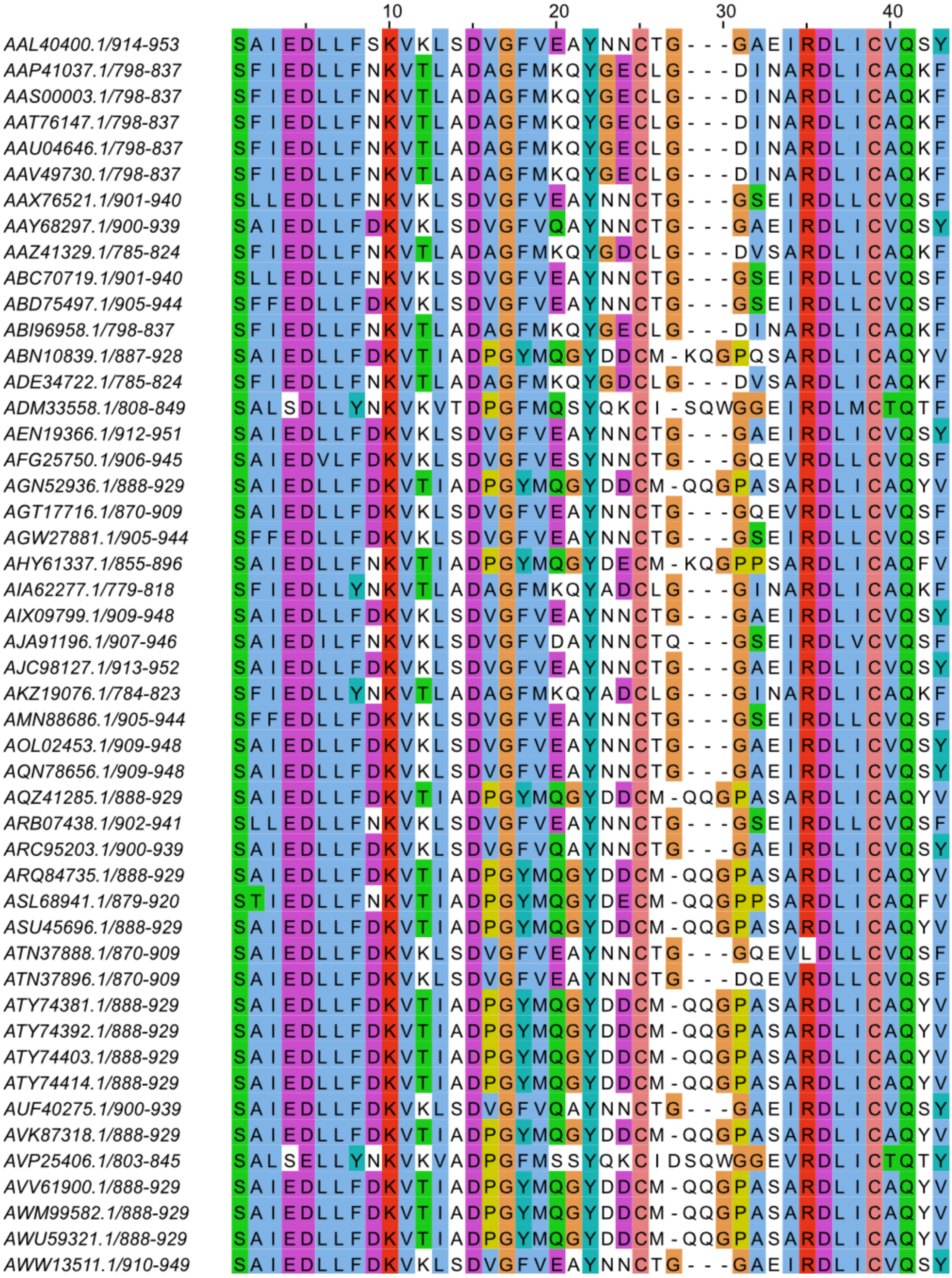

10
20
30
40
AAL40400.1/914-953 SAIEDLLFSKVKLSDVGFVEAYNNCTG---GAEIRDLICVQSY
AAP41037.1/798-837 SFIEDLLFNKVTLADAGFMKQYGECLG---DINARDLICAQKF
AAS00003.1/798-837 SFIEDLLFNKVTLADAGFMKQYGECLG---DINARDLICAQKF
AAT76147.1/798-837 SFIEDLLFNKVTLADAGFMKQYGECLG---DINARDLICAQKF
AAU04646.1/798-837 SFIEDLLFNKVTLADAGFMKQYGECLG---DINARDLICAQKF
AAV49730.1/798-837 SFIEDLLFNKVTLADAGFMKQYGECLG---DINARDLICAQKF
AAX76521.1/901-940 SLLEDLLFNKVKLSDVGFVEAYNNCTG---GSEIRDLLCVQSF
AAY68297.1/900-939 SAIEDLLFDKVKLSDVGFVQAYNNCTG---GAEIRDLICVQSY
AAZ41329.1/785-824 SFIEDLLFNKVTLADAGFMKQYGDCLG---DVSARDLICAQKF
ABC70719.1/901-940 SLLEDLLFNKVKLSDVGFVEAYNNCTG---GSEIRDLLCVQSF
ABD75497.1/905-944 SFFEDLLFDKVKLSDVGFVEAYNNCTG---GSEIRDLLCVQSF
ABI96958.1/798-837 SFIEDLLFNKVTLADAGFMKQYGECLG---DINARDLICAQKF
ABN10839.1/887-928 SAIEDLLFDKVTIADPGYMQGYDDCM-KQGPQSARDLICAQYV
ADE34722.1/785-824 SFIEDLLFNKVTLADAGFMKQYGDCLG---DVSARDLICAQKF
ADM33558.1/808-849 SALSDLLYNKVKVTDPGFMQSYQKCI-SQWGGEIRDLMCTQTF
AEN19366.1/912-951 SAIEDLLFDKVKLSDVGFVEAYNNCTG---GAEIRDLICVQSY
AFG25750.1/906-945 SAIEDVLFDKVKLSDVGFVESYNNCTG---GQEVRDLLCVQSF
AGN52936.1/888-929 SAIEDLLFDKVTIADPGYMQGYDDCM-QQGPASARDLICAQYV
AGT17716.1/870-909 SAIEDLLFDKVKLSDVGFVEAYNNCTG---GQEVRDLLCVQSF
AGW27881.1/905-944 SFFEDLLFDKVKLSDVGFVEAYNNCTG---GSEIRDLLCVQSF
AHY61337.1/855-896 SAIEDLLFNKVTIADPGYMQGYDECM-KQGPPSARDLICAQFV
AIA62277.1/779-818 SFIEDLLYNKVTLADAGFMKQYADCLG---GINARDLICAQKF
AIX09799.1/909-948 SAIEDLLFDKVKLSDVGFVEAYNNCTG---GAEIRDLICVQSY
AJA91196.1/907-946 SAIEDILFNKVKLSDVGFVDAYNNCTQ---GSEIRDLVCVQSF
AJC98127.1/913-952 SAIEDLLFDKVKLSDVGFVEAYNNCTG---GAEIRDLICVQSY
AKZ19076.1/784-823 SFIEDLLYNKVTLADAGFMKQYADCLG---GINARDLICAQKF
AMN88686.1/905-944 SFFEDLLFDKVKLSDVGFVEAYNNCTG---GSEIRDLLCVQSF
AOL02453.1/909-948 SAIEDLLFDKVKLSDVGFVEAYNNCTG---GAEIRDLICVQSY
AQN78656.1/909-948 SAIEDLLFDKVKLSDVGFVEAYNNCTG---GAEIRDLICVQSY
AQZ41285.1/888-929 SAIEDLLFDKVTIADPGYMQGYDDCM-QQGPASARDLICAQYV
ARB07438.1/902-941 SLLEDLLFNKVKLSDVGFVEAYNNCTG---GSEIRDLLCVQSF
ARC95203.1/900-939 SAIEDLLFDKVKLSDVGFVQAYNNCTG---GAEIRDLICVQSY
ARQ84735.1/888-929 SAIEDLLFDKVTIADPGYMQGYDDCM-QQGPASARDLICAQYV
ASL68941.1/879-920 STIEDLLFNKVTIADPGYMQGYDECM-QQGPPSARDLICAQFV
ASU45696.1/888-929 SAIEDLLFDKVTIADPGYMQGYDDCM-QQGPASARDLICAQYV
ATN37888.1/870-909 SAIEDLLFDKVKLSDVGFVEAYNNCTG---GQEVLDLLCVQSF
ATN37896.1/870-909 SAIEDLLFDKVKLSDVGFVEAYNNCTG---DQEVRDLLCVQSF
ATY74381.1/888-929 SAIEDLLFDKVTIADPGYMQGYDDCM-QQGPASARDLICAQYV
ATY74392.1/888-929 SAIEDLLFDKVTIADPGYMQGYDDCM-QQGPASARDLICAQYV
ATY74403.1/888-929 SAIEDLLFDKVTIADPGYMQGYDDCM-QQGPASARDLICAQYV
ATY74414.1/888-929 SAIEDLLFDKVTIADPGYMQGYDDCM-QQGPASARDLICAQYV
AUF40275.1/900-939 SAIEDLLFDKVKLSDVGFVQAYNNCTG---GAEIRDLICVQSY
AVK87318.1/888-929 SAIEDLLFDKVTIADPGYMQGYDDCM-QQGPASARDLICAQYV
AVP25406.1/803-845 SALSELLYNKVKVADPGFMSSYQKCIDSQWGGEVRDLICTQTY
AVV61900.1/888-929 SAIEDLLFDKVTIADPGYMQGYDDCM-QQGPASARDLICAQYV
AWM99582.1/888-929 SAIEDLLFDKVTIADPGYMQGYDDCM-QQGPASARDLICAQYV
AWU59321.1/888-929 SAIEDLLFDKVTIADPGYMQGYDDCM-QQGPASARDLICAQYV
AWW13511.1/910-949 SAIEDLLFDKVKLSDVGFVEAYNNCTG---GAEIRDLICVQSY

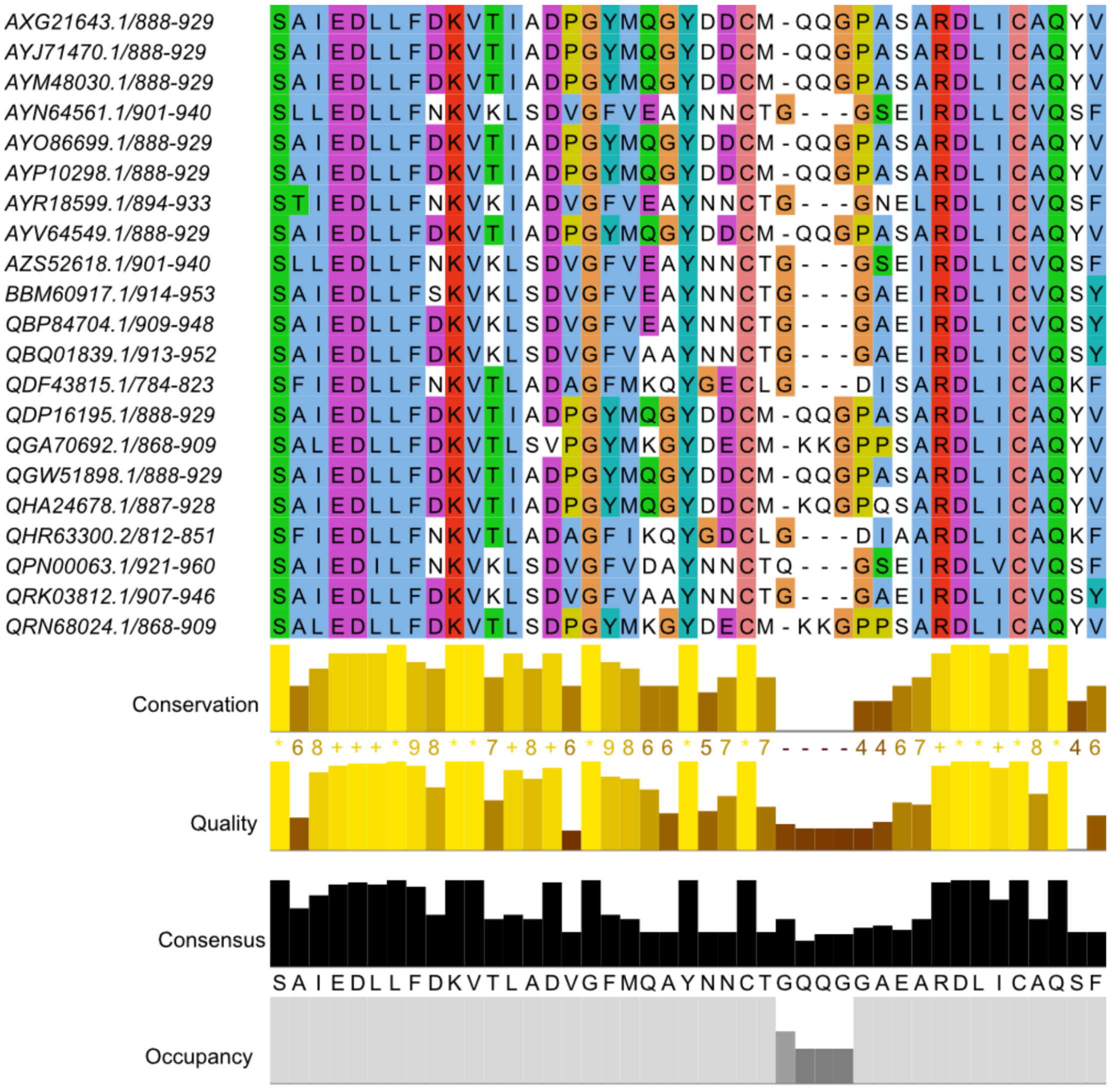


Figure S1: **Alignment of the FP region of betacoronaviruses.** Complete S protein sequences of the genus betacoronavirus were selected from the NCBI virus database[40] and aligned using Clustal Omega.[22] Only the FP region is shown. The fully conserved cysteines are at positions 25 and 39, respectively. Multiple entries from the same species were deleted. Coloring according to ClustalX.

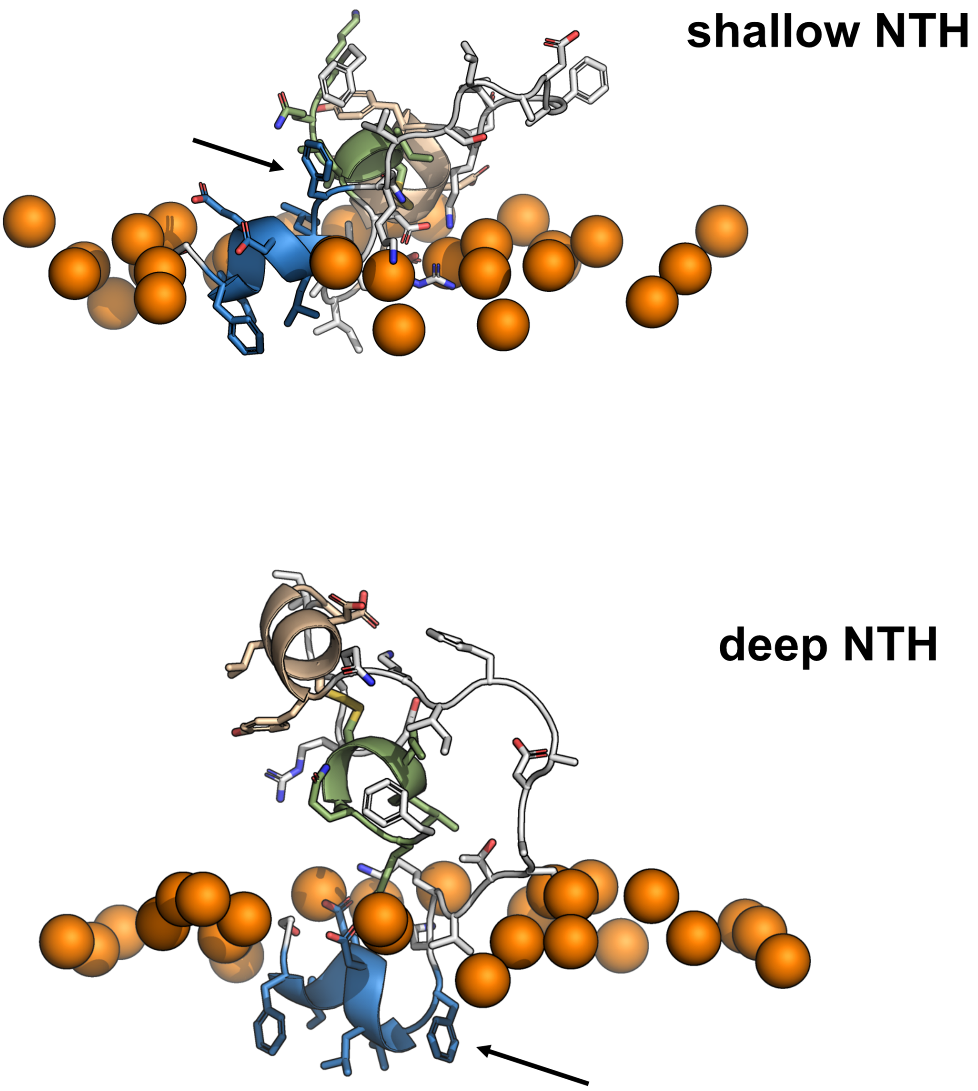


Figure S2: **Deep and shallow NTH insertion.** Arrows indicate the F823 position not inserted (top) and inserted (bottom) into the membrane interface. Colors as in Figure 1 (NTH: blue, AH2: beige, CTH: green). The upper membrane boundary is indicated by phosphate headgroups of nearby lipids (orange spheres).

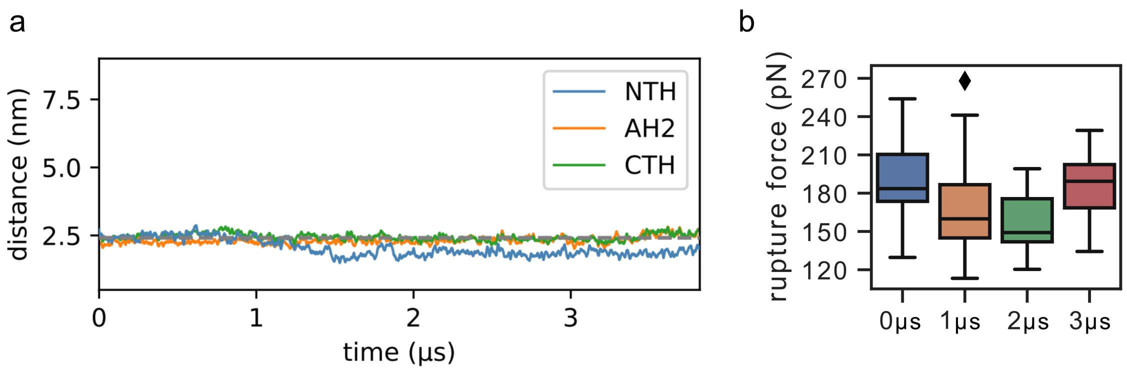


Figure S3: **Simulation starting with all three helices bound and adjusted outer-leaflet density.** (a) Distances of the centers of mass of the three helices and the center of mass of the membrane in which the lipid density of the outer leaflet was adjusted to accommodate the bound FP. The average phosphate position of the outer leaflet is indicated by a gray dotted line. (b) Box-plots showing the peak rupture forces of 20 replica pulling simulations each for pulling simulations started with structures of the simulation taken at 0, 1, 2, and 3 $\mu$s (box: interquartile range; median: horizontal line; diamond: outlier).

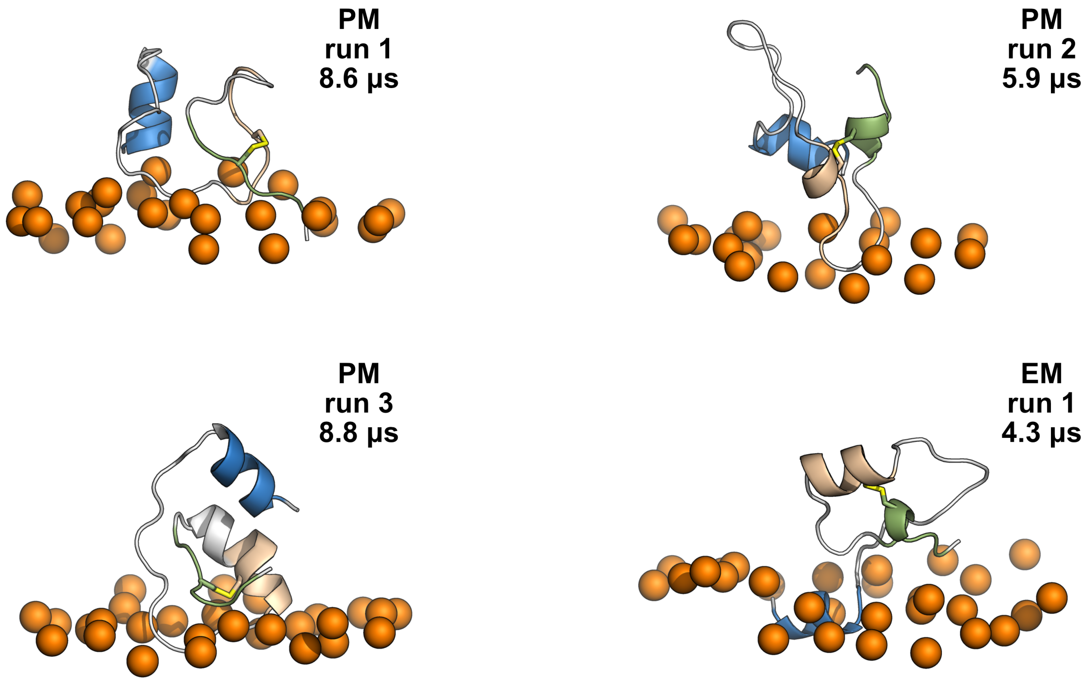


Figure S4: **Snapshots of MD simulations in which the AH2 and CTH partially unfolded to form flexible membrane bound structures.** (Left) FP interactions with the mimetic of the outer plasma membrane (PM). (Right) FP interactions with the mimetic of the endosomal membrane (EM). Time points are indicated. Colors as in Figure 1 (NTH: blue, AH2: beige, CTH: green). The upper membrane boundary is indicated by phosphate headgroups of nearby lipids (orange spheres).

### Videos

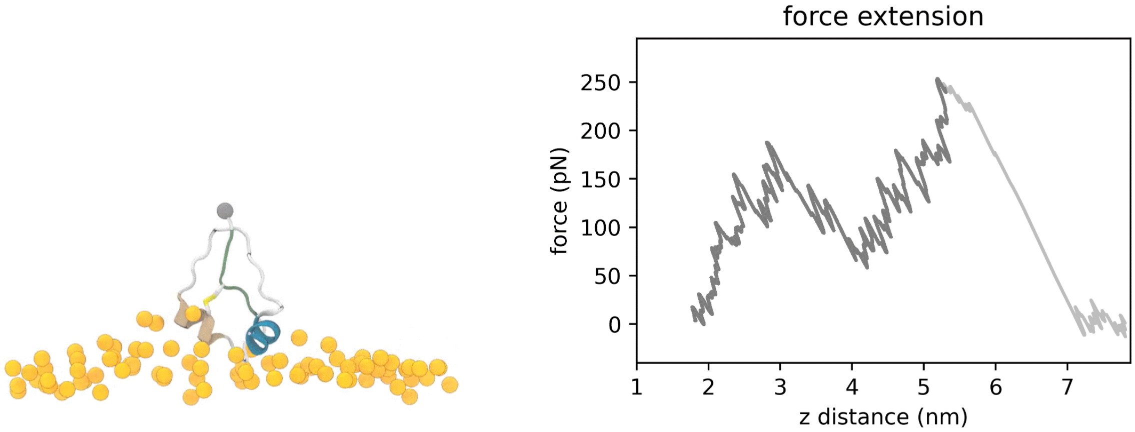


Figure S5: **Movie of FP pulling.** (Left) Trajectory of the pulling simulation that resulted in the highest rupture force of about 250 pN. The FP is shown in the same colors as used in Figure 1. The gray sphere indicates the C-terminus, to which the pulling force is applied. Lipid headgroup phosphates are shown as yellow spheres. (Right) Force extension curve corresponding to the trajectory. X-axis shows the distance of the C-terminal carbon (gray sphere) to the center of mass of the membrane. Y-axis shows the force acting on the C-terminal carbon.
